# Anabolic phenotype in cartilage-specific Mitogen-inducible gene-6 knockout mice is independent of Transforming growth factor-*α*

**DOI:** 10.1101/2023.01.31.526523

**Authors:** Ermina Hadzic, Bethia To, Michael A. Pest, Ling Qin, Frank Beier

**Affiliations:** Department of Physiology and Pharmacology, Schulich School of Medicine & Dentistry, Western University, ON, Canada; Department of Physiology and Pharmacology, Collaborative Specialization in Musculoskeletal Health Research, Western University, ON, Canada; Bone and Joint Institute, Western University, ON, Canada; Department of Orthopaedic Surgery, Perelman School of Medicine, University of Pennsylvania, PA, USA

## Abstract

**Background/Objective:** Osteoarthritis (OA) is a whole joint disorder with no disease modifying treatment currently available. The Epidermal Growth Factor Receptor (EGFR) signaling pathway plays an important role in cartilage/bone development and its ligand transforming growth factor-*α* (TGl·*α*) is upregulated in OA. In contrast, Mitogen-inducible gene 6 (Mig6) is a negative regulator of EGFR, and our studies demonstrate that cartilage-specific Mig-6 deletion results in anabolic effects on cartilage and formation of chondro-osseus nodules (CON). We aimed to attenuate EGFR signaling by inhibiting TGF*α* production in cartilage-specific Mig6 deficient mice, to test whether this would prevent the formation of CONs.

**Methods:** We generated double knockout mice by crossing cartilage-specific *Mig-6^fl/fl^Col2a1-Cre^+/-^* and whole-body *Tgfa^+/-^* mice to generate experimental and control wild-type mice. Knee and elbow sections were stained with toluidine blue to examine articular cartilage thickness and cell density, and tartrate-resistant acid phosphatase for osteoclast activity. Additionally, immunohistochemistry was completed to analyze phospho-EGFR and SOX9.

**Results:** Mig-6 deficient mice display cartilage thickening and CONs at 12 weeks in both the elbow and knee joints, which is independent of TGF*α* ligand presence. Similarly, articular cartilage cell density is increased in *Mig-6^fl/fl^Col2a1-Cre^+/-^Tgfa^-/-^* and *Mig-6^fl/fl^Col2al-Cre^+/-^* mice, but not *Tgfa^-/-^* mice, and displays increased SOX9 and phospho-EGFR staining.

**Conclusion:** The articular cartilage displays increased thickness, increased cell density, and CON formation independent of the presence of TGF*α*, suggesting the anabolic phenotype in the Mig6-deficient mice is independent of TGF*α*/EGFR binding. The anabolic phenotype may be due to an alternative EGFR ligand activation, or perhaps due to other non-EGFR specific mechanism. More research is required to elucidate the exact pathway responsible for the anabolic effects.

## 1. Introduction

Osteoarthritis (OA) is the most common type of arthritis, defined as a whole joint disorder characterized by cartilage degeneration, subchondral bone remodeling, osteophyte formation, and synovitis, with complex tissue cross-talk [1, 2]. The multifactorial nature of OA has proven to be a challenge in the effort to identify therapeutic targets and there is currently no disease modifying OA drug (DMOAD) to stop, slow, or reverse OA progression.

The Epidermal Growth Factor Receptor (EGFR) signaling pathway plays an important role in cartilage/bone development and homeostasis as it leads to the downstream activation of many important pathways such as MAPK/ERK, PI3K/AKT, and JAK/STAT, causing cell proliferation, differentiation, and survival [3]. The importance of EGFR cannot be overstated, as almost all cell types express EGFR family members and the knockout of any one of them is lethal in mice [4]. Interestingly, EGFR signaling has been reported to both attenuate and aggravate OA, demonstrating very context-dependent activity with factors such as disease stage, age, sex, and ligand-receptor combinations affecting outcomes [5].

Transforming growth factor-alpha (TGF*α*), an EGFR ligand, is increased in the synovial membrane and synovial fluid of OA patients [6]. Indeed, TGF*α* expression is increased 4-fold in rat chondrocytes following surgical induction of OA [7]. Our group previously demonstrated catabolic effects of TGF*α* in *ex vivo* rat osteochondral explants, including degradation of ECM proteins, loss of chondrocyte phenotype, chondrocyte clustering, and increased levels of the OA marker MMP-13 [8, 9]. Indeed, pharmacological inhibition of EGFR slows OA progression in a rat model of OA [10], and mice deficient for TGF*α* are protected from post-traumatic OA, at least at young age [11.

EGFR signaling is regulated by mitogen inducible gene 6 (Mig6), a scaffold protein that down-regulates EGFR signaling [12]. It is also referred to as Gene 33, ErbB receptor feedback inhibitor 1 (ERRFI1), or receptor-associated late transducer (RALT), although it will be referred to as Mig6 for this paper for simplicity. Mig6 plays an important role in joint homeostasis; in fact, whole-body *Mig-6* deficient mice develop joint deformities and most die within 6 months due to temporomandibular joint ankylosis [13]. Using the *Col2a1 Cre* system to specifically delete *Mig-6* from mouse chondrocytes provides improved survival compared to global knockout, and the mutant mice exhibit increased cartilage thickness. These mice also display severe osteophyte-like chondro-osseus nodules (CON) in the knee and spine, severely impairing ambulation [14, 15].

In the present study, we aim to attenuate EGFR signaling by inhibiting TGF*α* production in cartilage-specific *Mig-6* deficient mice. We hypothesized that eliminating one major EGFR ligand would reduce the effect of Mig6 deficiency, possibly preventing the formation of CONs in mice with an anabolic cartilage phenotype.

## 2. Materials and Methods

### 2.1 Animals and surgery

Mice deficient in TGF*α* were developed using embryonic stem cell gene targeting via phosphoglycerate kinase (PGK)-*neo* expression cassette by Mann and colleagues [16, 11]. Cartilage-specific Mig6 deficient mice were established by breeding *Mig-6^fl/fl^* mice [17] with *Col2a1Cre* mice [18]. We generated double knockout mice by crossing cartilage-specific *Mig-6^fl/fl^Col2a1-Cre^+/-^* and whole-body *Tgfa^-/-^* mice to generate experimental (*Mig-d^fl/fl^Col2a1-Cre^+/-^ Tgfa^-/-^,Mig-6^fl/fl^Col2a1-Cre^+/-^ Tgfa^+/+^, Mig-6^fl/fl^Col2a1-Cre^-/-^ Tgfa^-/-^*) and control wild-type mice (*Mig-6^fl/fl^Col2a1-Cre^-/-^ Tgfa^+/+^*). Henceforth, double-knockout mice are referenced as *Mig-6^fl/fl^Col2a1-Cre^-/-^ Tgfa^-/-^*, Mig6 deficient mice as *Mig-6^fl/fl^Col2a1-Cre^+/-^*, and TGF*α* deficient mice as *Tgfa^-/-^*, Male mice from each group were selected for further experimentation, using age-match litters to achieve N = 5. Mice were bred in-house and maintained in a temperature- and humidity-controlled room with water and standard chow freely available. All mice were genotyped using extracted DNA from ear biopsies at 21 days of age. Mice were euthanized by CO_2_ asphyxiation at 12-weeks of age. All animal experiments were in accordance with the Canadian Council on Animal Care guidelines and were approved by the Animal Use Subcommittee at Western University (2019-035).

### 2.2 Histological Assessment and Immunohistochemistry

Knee and elbow joints were dissected and fixed in 4% paraformaldehyde at 4 °C overnight. Joint decalcification with 5% ethylenediaminetetraacetic acid (EDTA) in phosphate buffered saline (PBS) at 7.0 pH was carried out for 11 days at room temperature. Following processing and paraffin embedding, joints were sagitally or frontally sectioned at 5 μm. Samples from the left knee and elbow were stained in 0.04% Toluidine Blue in 0.2M acetate buffer, pH 4.0 to assess glycosaminoglycan content and general histology.

Right elbow samples were dewaxed using xylene and rehydrated using a graded series of ethanol washes, then incubated in 3% hydrogen peroxide in methanol for 15 minutes to remove endogenous peroxidase activity. 0/1% Triton X-100 was used for antigen retrieval and then blocked with 5% serum in PBS. Sections were then stained overnight at 4°C using primary antibody for SOX9 (R&D Systems) or phosphor-EGFR (phosphoTyr-1173; Cell Signaling Technology), and incubated with secondary antibody conjugated to horseradish peroxidase

(Santa Cruz Biotechnology). Sections were exposed to DAB+ chromogen (Dako Canada) and counterstained with 0.5% methyl green in 0.1M sodium acetate buffer 4.2 pH. Tartrate-resistant acid phosphatase (TRAP; Sigma Canada) staining was also performed to assess osteoclast activity. TRAP staining was done in accordance to manufacturers instructions. Images for all stained slides were taken on a Leica DFC295 digital camera affixed to a Leica DM1000 microscope.

### 2.3 Articular Cartilage Measurement

Sagittal knee and elbow sections were blinded and randomized for analysis. The articular cartilage thickness was measured at three evenly spaced points and averaged. Measurements were done on a minimum of three sections per joint per animal, with an average of 50 μm between sections. The Leica Application Suite software (v3.8.0) was used.

### 2.4 Cell Density

Cell density was calculated within central regions (200 μm wide x 70 μm tall) on the articular cartilage of the humerus. Lacunae with nuclear staining were counted as cells using the Leica Application Suite software (v3.8.0) on a minimum of three sections per animal. Sections were blinded and randomized for analysis.

### 2.5 Statistical Analysis

Statistical analysis was performed in GraphPad Prism v.9.5.0. Normality was assessed via the D’Agostino & Pearson test. A one-way analysis of variance (ANOVA) with Tukey’s multiple comparison test was run for articular cartilage thickness and cell density. Data is presented as mean with 95% CI. Data was considered statistically significant when p < 0.05.

## 3. Results

### 3.1 Anabolic articular cartilage phenotype in Mig6 deficient mice

All genotypes were confirmed through PCR, and phospho-EGFR immunohistochemistry was performed to demonstrate EGFR activity. As expected, phospho-EGFR staining was increased in Mig6-deficient mice compared to wild-type mice (**Fig. 1**). TGF*α*-deficient mice had fewer positive cells compared to the wild-type and Mig6 deficient groups. Double knockout mice appeared to show a similar number of positive cells as *Mig-6^fl/fl^Col2a1-Cre^+/-^* mice (**Fig. 1**). The thickness of articular cartilage was significantly increased in the femur (**Fig. 2 A, E**), tibia (**Fig. 2 B, E**), humerus (**Fig. 2 C**), and ulna (**Fig. 2 D**) of cartilage-specific Mig6 deficient mice, regardless of the presence of TGF*α*. There was no evidence of changes in articular cartilage thickness in *Tgfa^-/-^* mice compared to wild-type mice in knee or elbow joints. Interestingly, there was no difference in articular cartilage thickness between the double knockout and Mig6 deficient mice (femur mean difference = −20.99, 95% CI [−47.65, 5.67]).

**Figure 1.**
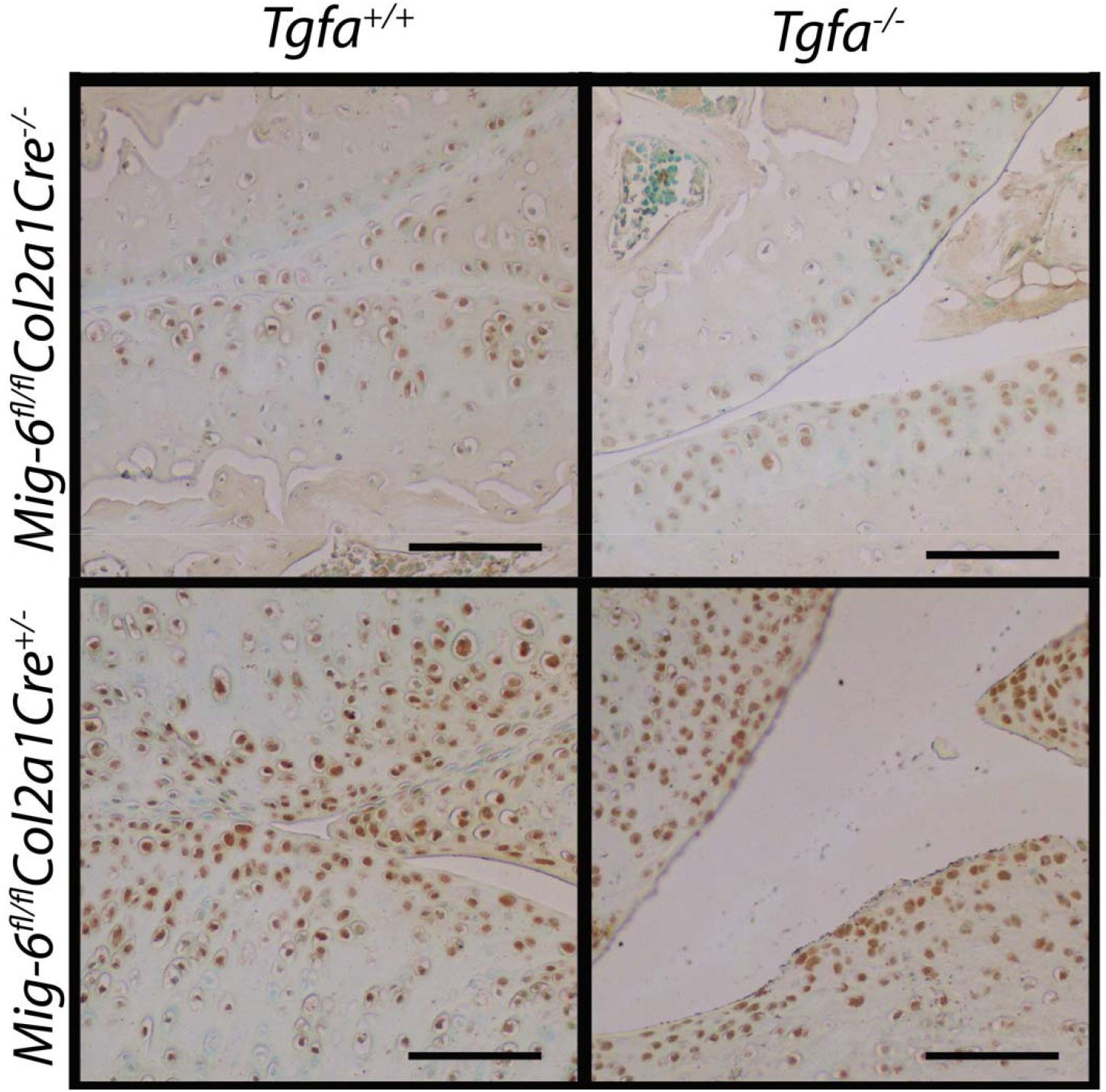
phospho-EGFR activity is upregulated in Mig6 deficient mice. Brown staining indicates positive phospho-EGFR staining, which appears more intense and present in more cells in Mig6 deficient mice, with opposite effects in *Tgfa^-/-^* mice. The *Mig-6^fl/fl^Col2a1-Cre^+/-^ Tgfa^-/-^* mice seem to have a similar amount of staining to the *Mig-6^fl/fl^Col2a1-Cre^+/-^* mice. Images represent sagittal right elbow sections, counterstained in methyl green. Scale bars = 100 μm. N = 5.

**Figure 2.**
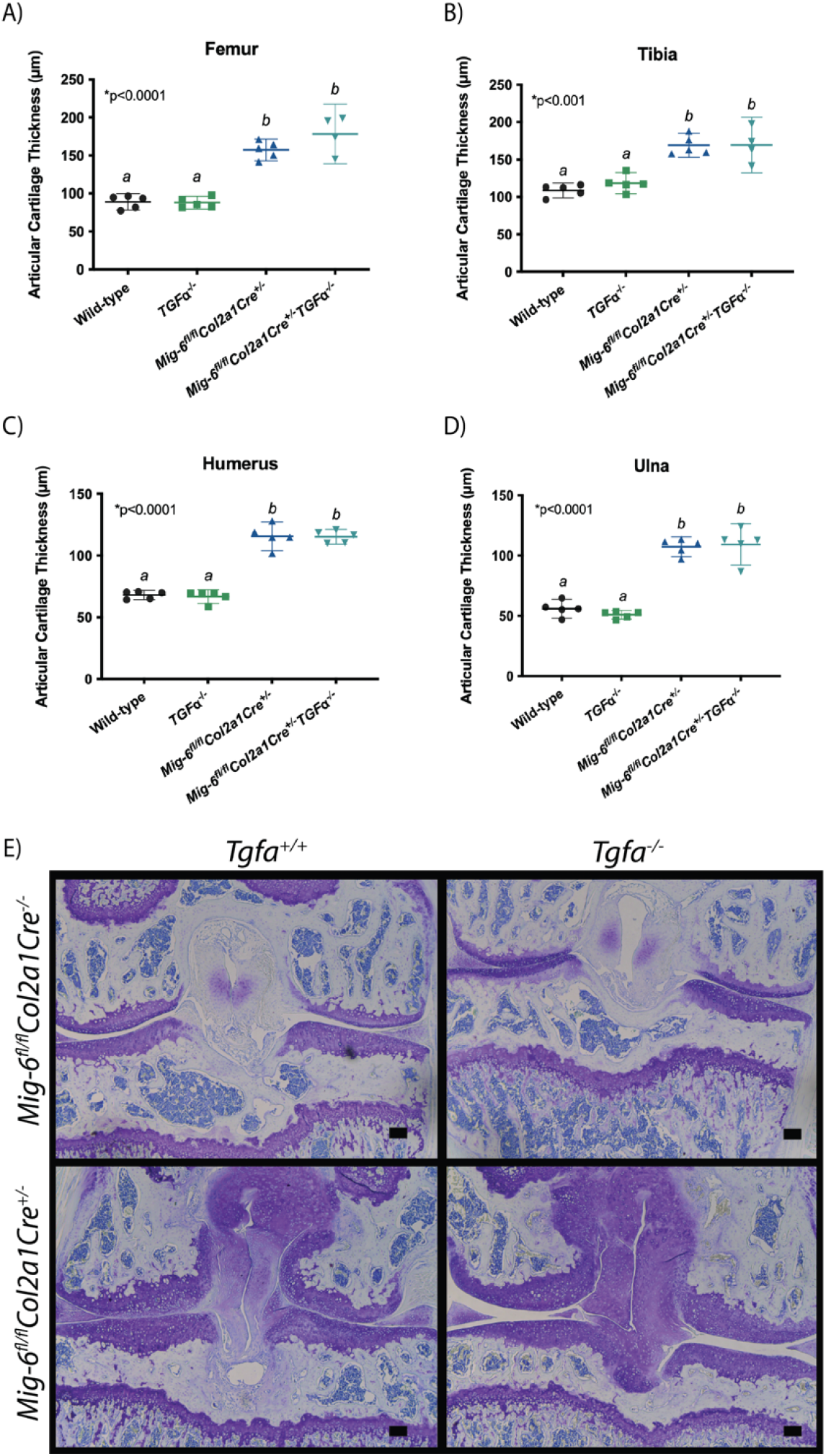
Mig6 deficient mice display articular cartilage thickening at 12 weeks, regardless of TGF*α*. Average articular cartilage thickness was calculated using three measurements at different locations on the articular cartilage using Leica Software. Average articular cartilage thickness was significantly increased in Mig6 deficient mice, regardless of TGF*α* presence in the (A, E) femur, (B, E) tibia, (C) humerus, and (D) ulna. Graphed data represents a non-parametric one-way ANOVA with Tukey’s multiple comparison test, where the individual mean and 95% CI is presented. (E) Representative images of toluidine blue stained knee sections. Scale bars = 100 μm. N = 5.

### 3.2 Cell density is not affected by TGF*α* deficiency

Mice with cartilage-specific deletion of Mig6 (with and without TGF*α* presence) were found to have a 1.5-fold increase in cell density in the cartilage of the humerus, compared to wild-type (*Mig-6^fl/fl^Col2a1-Cre^+/-^* mean difference = −21.75, 95% CI [−28.03, −15.47]; *Mig-6^fl/fl^Col2a1-Cre^+/-^ Tgfa^-/-^* mean difference = −22.69, 95% CI [−28.97, −16.41]) and TGF*α*-deficient mice (*Mig-6^fl/fl^Col2a1-Cre^+/-^* mean difference = −23.89, 95% CI [−30.17, −17.61]; *Mig-6^fl/fl^Col2a1-Cre^+/-^ Tgfa^-/-^* mean difference = −24.83, 95% CI [−31.11, −18.55]; **Fig. 3**). Similar to articular cartilage thickness, there was no difference between the double knockout and Mig6-deficient mice (mean difference = −0.941, 95% CI [−7.221, 5.339]). Immunohistochemistry staining for SOX9 was positive in all groups, although it appeared darker in the *Mig-6^fl/fl^Col2a1-Cre^+/-^ Tgfa^-/-^* and *Mig-6^fl/fl^Col2a1-Cre^+/-^* mice compared to the TGF*α*-deficient and wild type mice (**Fig. 4**).

**Figure 3.**
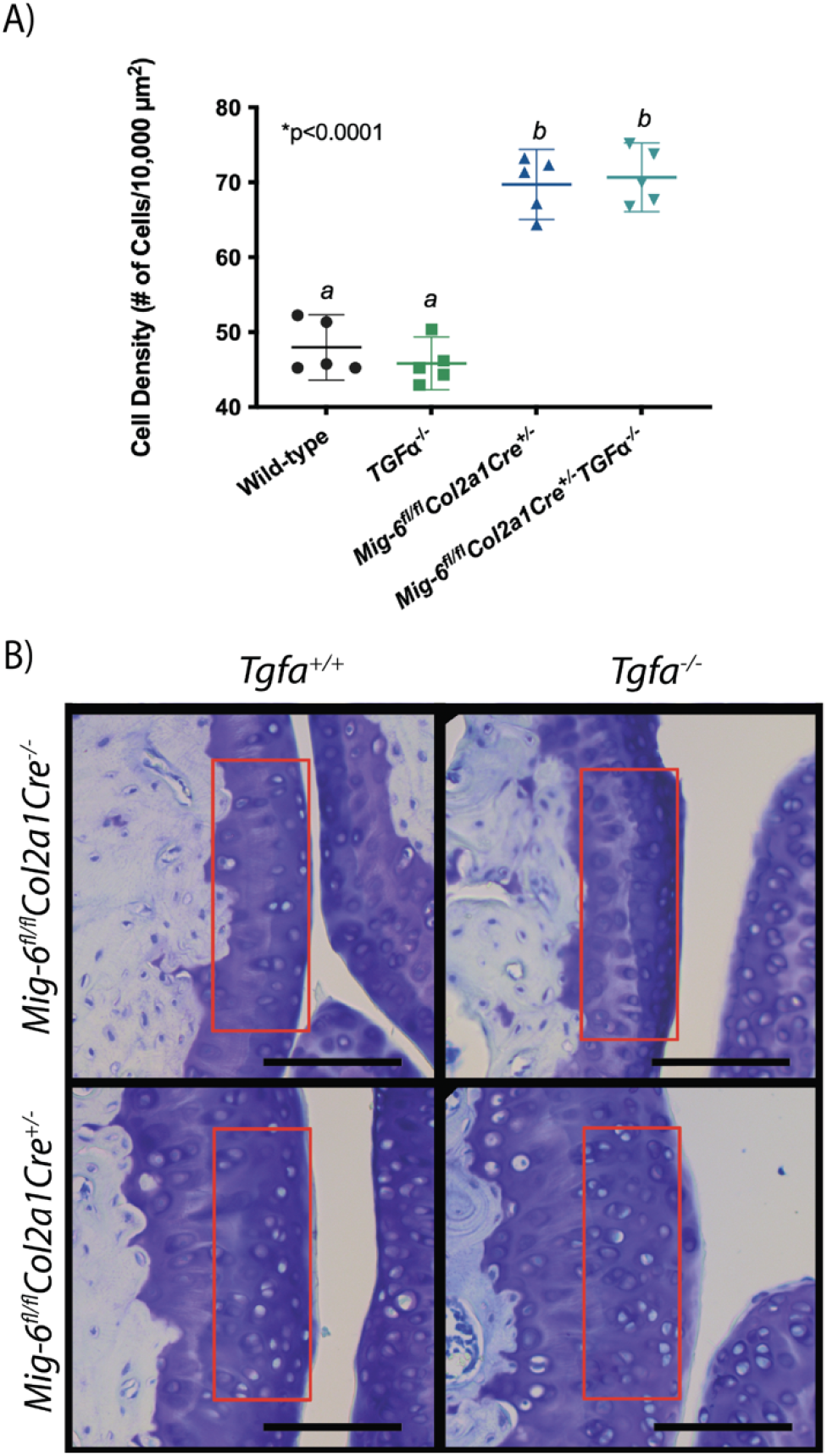
Cell density is not affected by TGF*α* deficiency. Cell density was calculated within the articular cartilage of toluidine blue stained elbow joints. Lacunae with nuclear staining was counted within a 70 μm x 200 μm region, indicated by the red box (B). Representative images of the elbow is presented. Scale bars = 100 μm. (A) Cell density is significantly increased in the humerus of Mig6 deficient mice, regardless of TGF*α* presence. Graphed data represents a non-parametric one-way ANOVA with Tukey’s multiple comparison test, where the individual mean and 95% CI is presented. N = 5.

**Figure 4.**
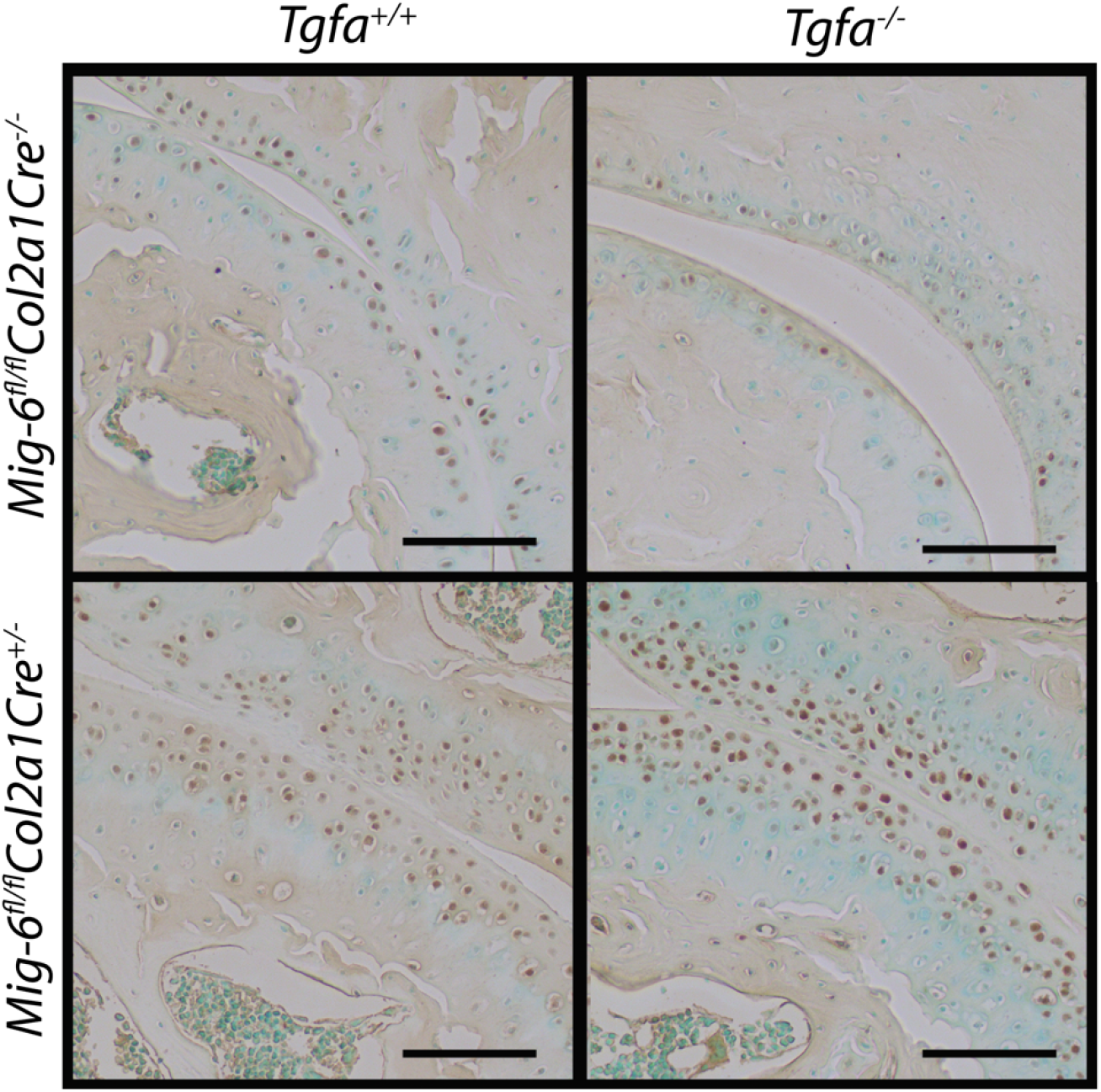
SOX9 staining is increased in Mig6 deficient groups. Brown staining indicates positive SRY-box transcription factor 9 (SOX9) nuclear staining, which is present in all groups. The *Mig-6^fl/fl^Col2a1-Cre^+/-^Tgfa^-/-^* mice and *Mig-6^fl/fl^Col2a1-Cre^+/-^* mice appear to have a darker stain than the *Tgfa^-/-^* and wild-type mice. All sections are of the sagittal right elbow, counterstained with methyl green. Scale bars = 100 μm. N = 5.

### 3.3 CONs develop in Mig6 deficient mice independent of TGF*α* ligand presence

Chondro-osseus nodules (CONs) around the articulating knee surface were present in double knockout (3 of n = 5) and cartilage-specific Mig6-deficient mice (2 of n = 5). CON development does not appear in wild-type or *Tgfa^-/-^* mice (**Fig. 5 A-B**). TRAP staining reveals an increase in osteoclast activity within CONs, which appears similar between the two Mig6-deficient groups (**Fig. 5 C**).

**Figure 5.**
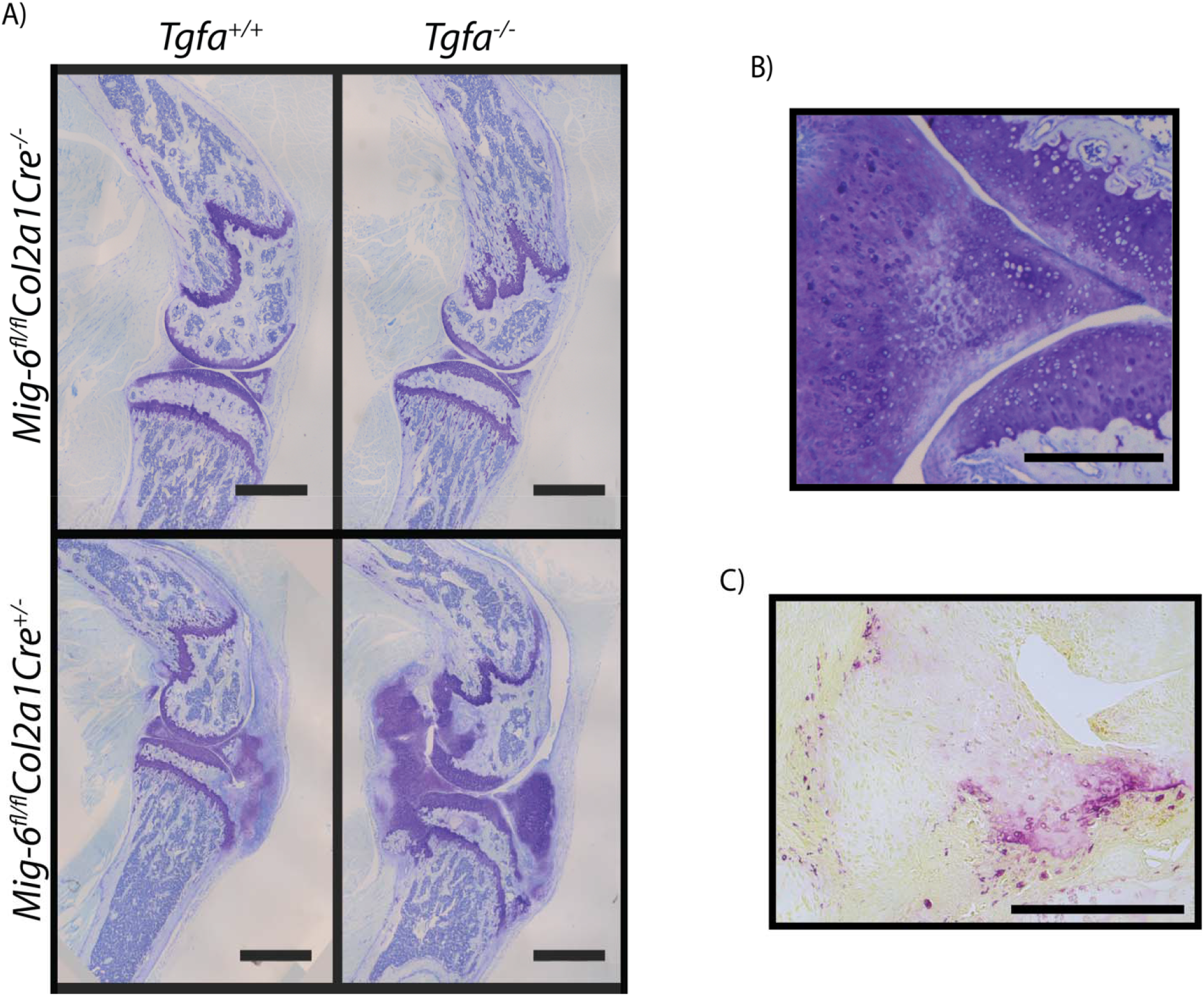
CONs develop in Mig6 deficient mice independent of TGF*α* ligand presence. (A) Toluidine blue stained sagittal knee sections of 12-week-old mice demonstrate CON development in Mig6 deficient mice, with or without TGF*α*. CON development does not appear in wild type or TGF*α* deficient mice. Images are representative of N = 2 for the *Mig-6^fl/fl^Col2a1-Cre^+/-^Tgfa^-/-^* mice and N = 3 for the remaining groups. Scale bars = 4 mm. (B) CONs have a greater dark purple proteoglycan staining in toluidine blue sections. Scale bar = 100 μm. (C) TRAP staining reveals an increase in osteoclast activity within CONs, which appears similar between groups. Scale bar = 50 μm.

## 4. Discussion

Regulation of the EGFR pathway via Mig6 signaling may be an interesting therapeutic target for OA patients, as Mig6 deficient mice develop an anabolic cartilage phenotype. In the present study, we aimed to assess the relationship TGF*α* plays in this phenotype, e.g. to examine whether absence of one major EGFR ligand would affect the anabolic effects of Mig6-deficiency. We noted that upon *Mig6* deletion, the articular cartilage displays increased thickness, cell density, and CON formation independent of TGF*α*, suggesting the anabolic phenotype in the Mig6 mice is independent of TGF*α*/EGFR binding. These data do not support our initial hypothesis.

Our results demonstrate an increase in articular cartilage thickness in both the knee and elbow joints of 12-week old *Mig-6^fl/fl^Col2a1-Cre^+/-^ Tgfa^-/-^* and *Mig-6^fl/fl^Col2a1-Cre^+/-^* mice, similar to previous studies. Pest et al. [12] reported a ~1.5-fold increase in knee articular cartilage thickness in *Mig-6^fl/fl^Col2a1-Cre^+/-^* mice, and we have similarly noted a ~1.6-fold increase. Our data on cellular density, SOX9, and EGFR staining is additionally similar to Pest et al. [12]. In contrast, we noted CON development in 2-3 animals with Mig6 deficiency (with or without TGF*α* presence) at 12-weeks of age, whereas Pest et al. [12] noted all Mig6 deficient mice developed CONs in at least 1 knee joint. This may be due to a sex difference, as our work was solely in male mice whereas Pest et al. [12] noted this occurrence in female mice. Moreover, we analyzed only one knee per animal. Unfortunately, the N = ≤ 3 prevents us from doing further analysis on those mice that did develop CONs.

The transcription factor SOX9 is a key marker of chondrocyte development and phenotype maintenance [19]. In our study, SOX9 antibody staining was increased in Mig6-deficient mice, which was expected due to the increase in articular cartilage thickness and cell density. Interestingly, the stain appears more intense in the double knockout mice compared to the Mig6-deficient mice. This finding is consistent with previous work demonstrating a reduction of SOX9 mRNA and protein expression following *in vitro* TGF*α* treatment [8]. We aim to quantify SOX9 (and phospho-EGFR) protein expression in these groups in future studies to validate the results.

We did not note a decrease in CON formation or cartilage thickening in the double knockout mice compared to *Mig6* KO mice, which we had expected. This may be due to a number of factors. Firstly, EGFR has multiple binding ligands so perhaps a ligand other than TGF*α* is responsible or compensates [3]. For example, heparin binding EGF-growth factor (HB-EGF) is an EGFR ligand that may be protective of surgically-induced OA in mice when it is overexpressed [20]. The relationship between EGFR and OA is very context dependent [5], and so exploring alternative ligand activation is a worthwhile future endeavor to understanding its role.

Other non-EGFR specific mechanisms may also be responsible for our findings. Mig6 is a negative regulatory of the c-Met receptor, which mediates cell proliferation and migration via hepatocyte growth factor (HGF) [21, 22]. Previous studies have demonstration HGF and c-Met expression by chondrocytes *in vivo*, both in healthy and OA tissue [23]. HGF and c-Met are produced by the calcified and deep zone chondrocytes [21]. The calcified cartilage and deep zone chondrocytes have a larger portion of c-Met receptors, compared to the mid and superficial zone chondrocytes [21]. Additionally, there is research pointing to a change in HGF/c-Met activity in synovial joints during OA [24]. It is therefore possible that our *Mig-6^fl/fl^Col2a1-Cre^+/-^* mice de-regulate c-Met signaling, which may contribute to the anabolic phenotype.

Overall, it appears that the anabolic phenotype found in Mig6 deficient mice is independent of TGF*α*/EGFR binding. There are a number of other ligand interactions our group will examine in the future to understand the mechanisms behind this anabolic phenotype.

## 5. Acknowledgements

The authors would like to thank all members of the Beier lab for ongoing support.

